# Cell rounding causes genomic instability by dissociation of single-stranded DNA-binding proteins

**DOI:** 10.1101/463653

**Authors:** Qian Guo, Xianglu Liao, Xingwu Wang, Ling Liu, Bao Song

## Abstract

Genomic instability can cause a wide range of diseases, including cancer and cellular senescence, which is also a major challenge in stem cell therapy. However, how a single event can cause extremely high levels of genomic instability remains unclear. Using our developed method, cell in situ electrophoresis (CISE), and models of normal, cancer, and embryonic stem cells, we found that cell rounding as a catastrophic source event ubiquitously observed in vivo and in vitro might lead to large-scale DNA deprotection, genomic instability, chromosomal shattering, cell heterogeneity, and senescent crisis by dissociation of single-stranded DNA-binding proteins (SSBs). Understanding the mechanism may facilitate the development of clinical strategies for cancer therapy, improve the safety of stem cell therapy, and prevent pathological aging.

## Main Text

DNA replication is a key process of cell proliferation involved in body growth, tissue renewal, and injury repair. The cascade of sequential biological events, i.e., cell attachment to the substrate, spreading, increasing nuclear size, and loosening of tightly packaged chromosomes into chromatin, is a prerequisite for DNA replication of normal somatic cells. Additionally, the increased nuclear size is essential for the conformational changes in chromatins and the assembly of replication complexes in which single-stranded DNA-binding proteins (SSBs) participate. These SSBs include replication protein A (RPA) and protection of telomere 1 (POT1), which are essential for protection of DNA from the formation of erroneous structures during DNA replication and for formation of complexes involved in DNA repair (1-7). Interruption of DNA replication by nonphysiological cell rounding, a type of global, fundamental event observed, for example, in inflammation (8) in vivo and cell culture in vitro, may become a source of stress for ongoing DNA replication. This type of cell rounding physically produces a cascade of releasing the puling force maintained by the cell-substrate adhesion, disturbing the balance of prestress force on the nucleus, making the compacting force generated by microtubules predominant, and finally reducing nuclear size (9). The microtubule-induced spatial-compaction reduction of nuclear size may alter the conformational states of both SSBs and chromatin, further resulting in abnormal changes in the binding states of SSBs, which are catastrophic to genomic stability (1, 2). Additionally, based on the observation that each human proliferating cell contains 40,000–80,000 origins of DNA replication (10) and may be at a different point in the phase of the cell cycle, one-time cell rounding may produce a type of microtubule-induced genomic instability (MIGI) by large-scale deprotection of DNA replication due to SSB unbinding. Understanding the global role of cell rounding in the MIGI may be helpful for cancer therapy, aging prevention, and safety of stem cell therapy. However, experimental confirmation of the theoretical hypothesis is a solid challenge because of the lack of methods to examine the binding states of SSBs under conditions in which cellular prestress remained intact. To this end, we developed a nondestructive approach to test and verify the mechanism, which may facilitate clinical applications.

To achieve this goal, we choose POT1, RPA70, and RPA32 as tested objectives; RPA14 was excluded owing to the lack of commercial availability of an anti-RPA14 antibody. Additionally, human fetal lung fibroblasts (HFLFs) and mouse embryonic fibroblasts (MEFs), HeLa cells, and mouse embryonic stem cells (mESCs) were chosen as models of differentiated normal somatic cells, tumor cells, and totipotent stem cells, respectively. To test the effects of cell rounding on nuclear reduction, all of the cells were canonically trypsinized for passage and experimental examination. To test the binding states of SSBs in a nondestructive manner, we developed a novel approach (see Supplemental Materials and Methods), called cell in situ electrophoresis (CISE; Fig. 1A), in which the unbound SSBs were separated from the integral cells. After CISE, the tested cells were pooled together, and the remaining SSBs in the cells, which represented the bound SSBs, were examined by flow cytometry (FCM). As the first step of using this combined method, we detected the binding states of RPA70, RPA32, and POT1 in HFLFs, HeLa cells, and mESCs. The results showed that the binding levels of the SSBs in rounding HFLFs were obviously lower (Fig. 1B), suggesting that the nuclear compaction caused by the cell rounding could make SSBs dissociated from their bound DNA. In contrast, no significant changes in the SSB binding states were observed in HeLa cells and mESCs compared with those in HFLFs (Fig. 1B). This result could be explained by the global nature of cancer and embryonic stem cells, which have relatively larger nuclei than the differentiated normal somatic cells (11-14) even after they become rounding. The changes in nuclear sizes before and after rounding are shown in Fig. 1C.

**Fig. 1.**
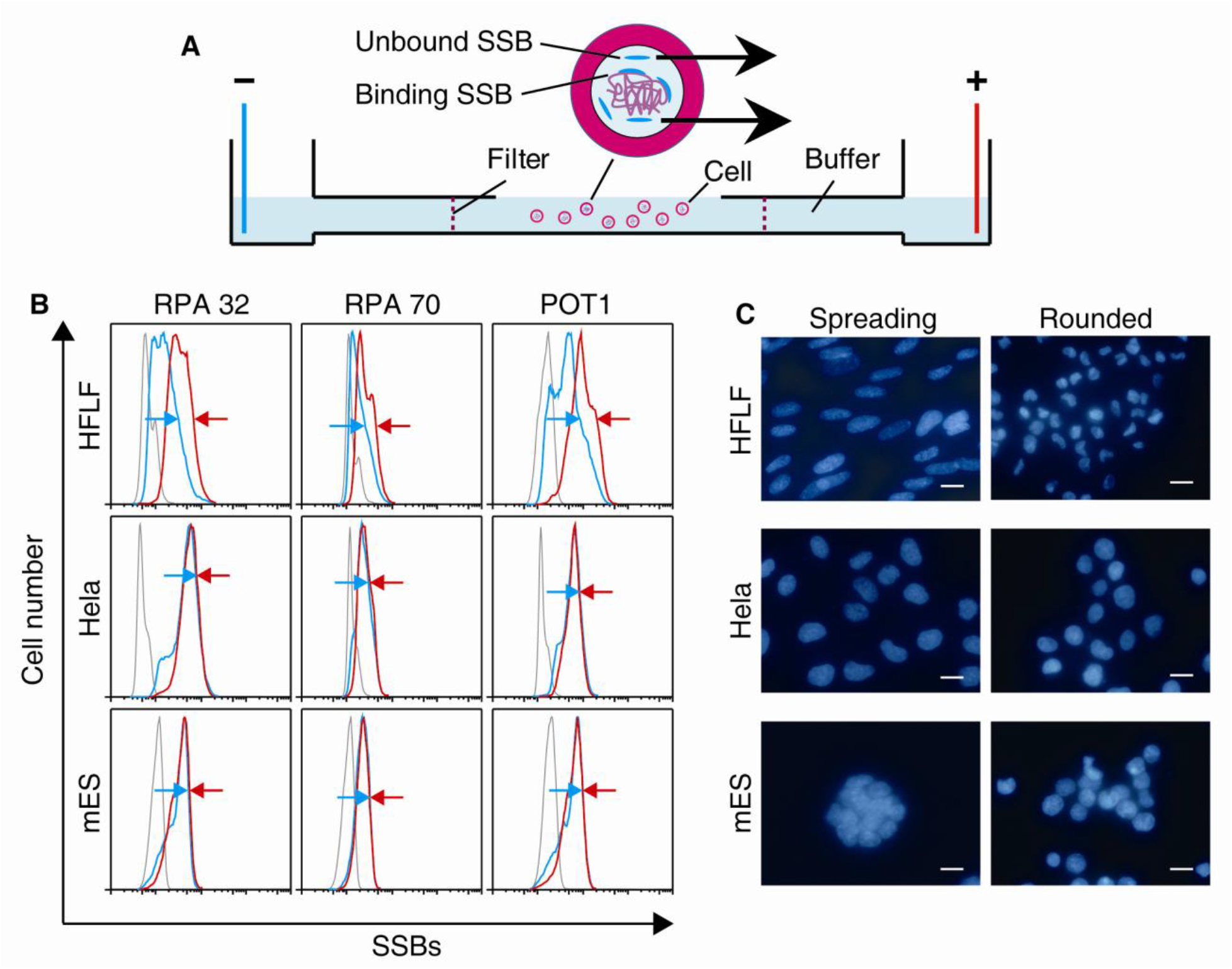
Changes in nuclear sizes and examination of SSB binding states through CISE. (**A**) Schematic diagram of the CISE. (**B**) Binding states of SSBs for rounded HFLFs, HeLa cells, and mESCs. Red line, no treatment; blue line, treatment with CISE; dark line, isotype control. The forward shift of back slopes of the peaks (blue) demonstrates the dissociated state of SSBs for HFLFs, which is in contrast to HeLa and mESCs. The gap between the paired arrows (green and red) indicates the obvious difference. (**C**) Changes in the nuclear sizes of HFLFs, HeLa cells, and mESCs. Scale bar, 15 μm.

To verify the source role of microtubules in generating compaction force of determining nuclear size and binding state of SSBs, we applied an anticancer drug, vincristine (VCR), as a tool to alter the functional states of microtubules. The effect of VCR on targeting microtubule dynamics can increases the nuclear size by disassembling microtubules (15). As anticipated, VCR-treated HFLFs remained enlarged nuclear size after rounding (Fig. 2A). This was consistent with the observation that the microtubule system plays a key role in controlling nuclear size (16, 17). More importantly, the changes of nuclear size also determined the corresponding changes of the binding states of the SSBs (Fig. 2B). However, the VCR treatment had no further significant effect of enlarging nuclear size and bound SSBs on mESCs (Fig. 2C and 2D), which remained similar to the cells without VCR treatment (Fig. 1B and 1C).

**Fig. 2.**
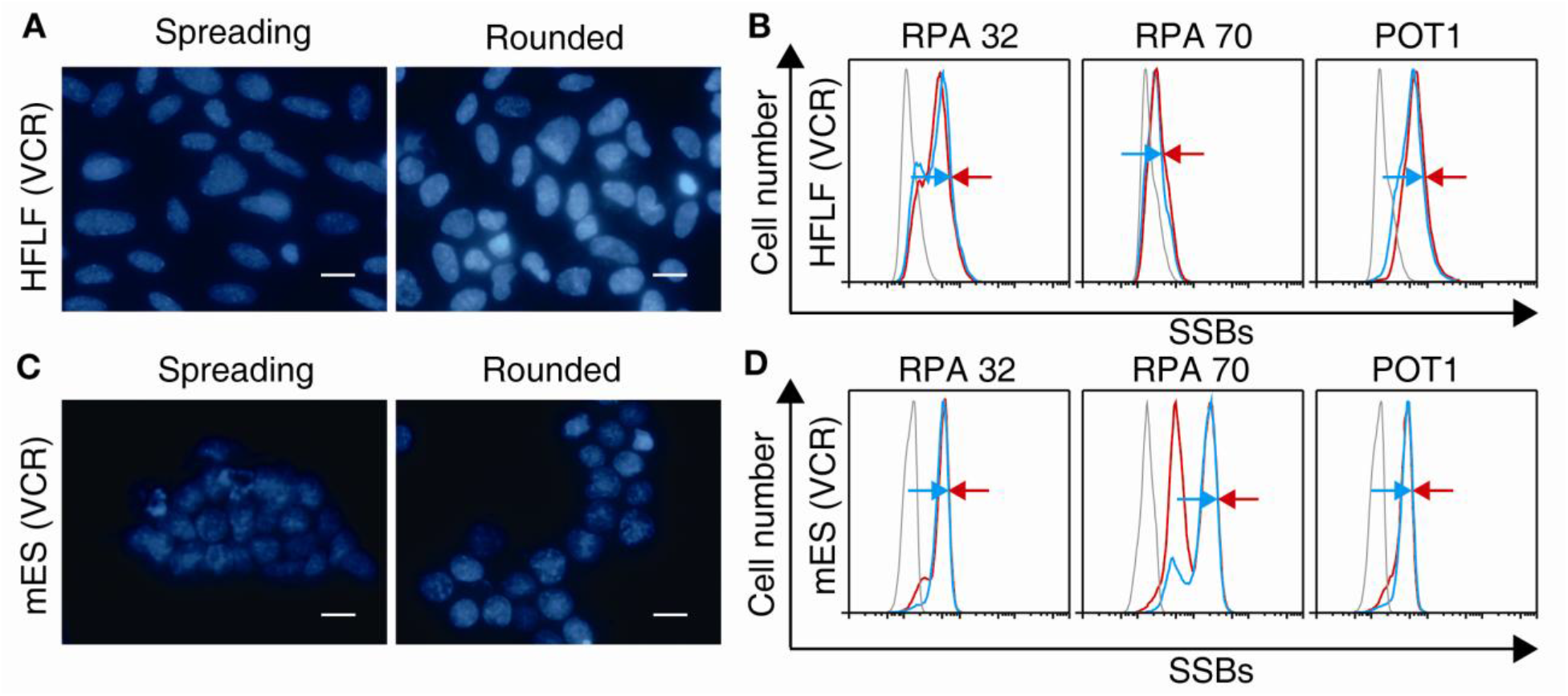
The role of microtubules in controlling nuclear size and binding states of SSBs. (**A**) Nuclear sizes of VCR-treated HFLFs before and after rounding. Scale bar, 15 μm. (**B**) SSB binding states of the rounded cells as in (A). Red line, no treatment; blue line, treatment with CISE; dark line, isotype control. The gap between the paired arrows (blue and red) shows the results of the drug treatment in the cells. (**C**) Nuclear sizes of VCR-treated mESCs before and after rounding. Scale bar, 15 μm. (**D**) SSB binding states of the rounded cells as in (C). Red line, no treatment; blue line, treatment with CISE; dark line, isotype control. The gap between the paired arrows (green and red) shows the results after drug treatment of the cells.

To further investigate the functional states of microtubules in differentiated and undifferentiated cells, we tested the effects of VCR on MEFs which are homologous with mESCs. The results showed that the MEFs with VCR treatment presented no significant nuclear compaction after rounding (Fig. 3A and 3B), which was similar to the effect of VCR on HFLFs (Fig. 2A and 2B). However,the effect of VCR on the nuclear enlargement and the binding state of SSBs could not be reflected in the undifferentiated cells, mESCs, due to their already possessed nature of no nuclear compaction after rounding (Fig. 2C and 2D). All of these suggest that the assembly of microtubules, as a modulator for stemness, may play a key role in determining the states of differentiation, because the qualitative abnormality of microtubules are deadly for mitotic cells. In addition, the state of microtubules as a key factor may also take part in forming the cascade of causality for genomic stability. Indeed, a recent study presented that HeLa cells typically exhibit high genomic stability (18), which may be explained by the above experimental results that the cells without nuclear compaction after rounding can prevent dissociation of SSBs and their bound DNA.

**Fig. 3.**
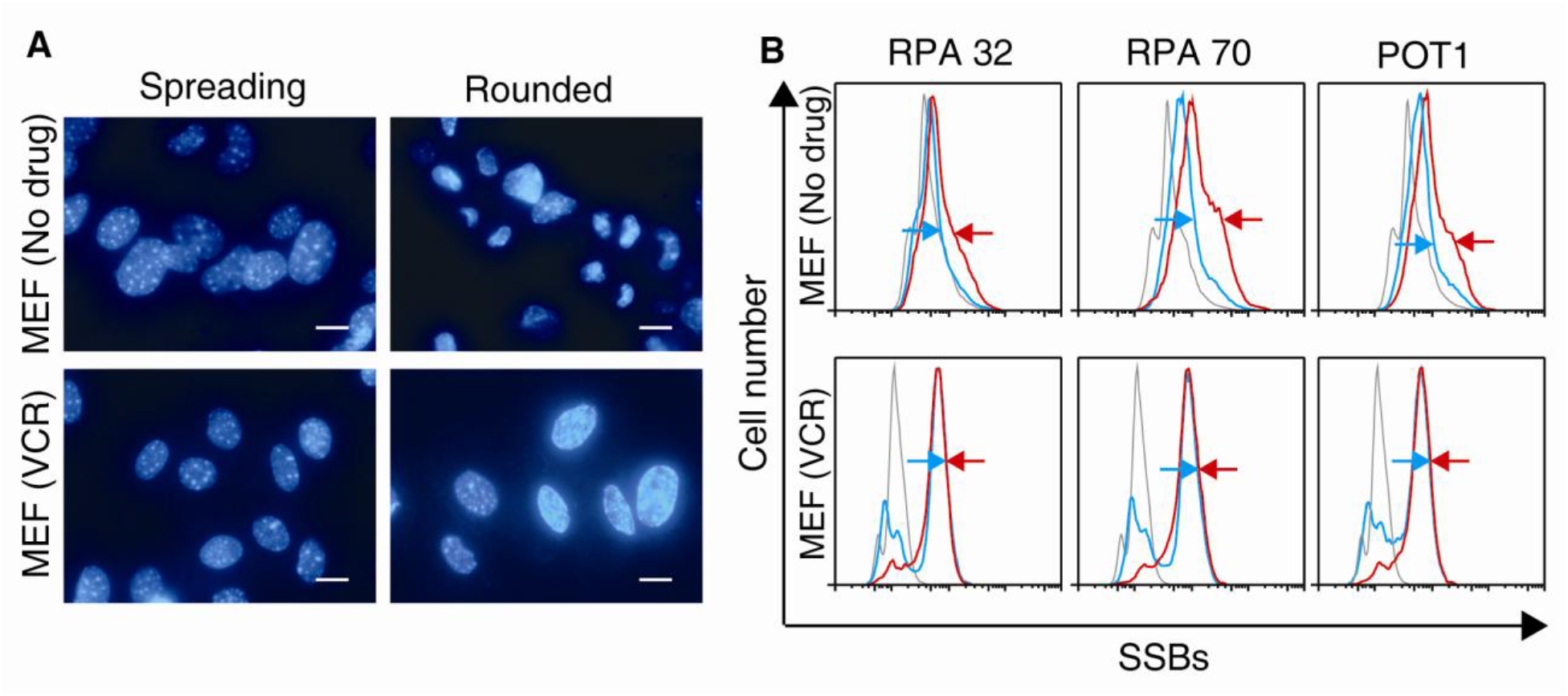
The role of microtubules in the homologous differentiated MEFs. (**A**) Nuclear sizes of untreated and VCR-treated MEFs before and after rounding. Scale bar, 15 μm. (**B**) SSB binding states of the cells as in (A). Red line, no treatment with CISE; blue line, treatment with CISE; dark line, isotype control. The gap between the paired arrows (blue and red) shows the different results.

Since SSB binding is a vital process for DNA replication and maintenance, loss of protection of POT1 causes immediate genome instability (1, 2). Therefore, events causing dissociation of SSBs in replicating cells may cause cellular genetic disorders and senescence. In order to examine the effects of such events, i.e., rounding, on cell replicative senescence, we applied a canonically experimental model designed by Hayflick to test cell senescence in vitro (19, 20), except that the passage time was modified to twice per day (Fig. 4A). The results demonstrated that the HFLFs became senescent and died after only about seven population doublings (PDs; Fig. 4A–C). This PD number is much shorter than that (~50 PDs) of the Hayflick limit (19, 20). Interestingly, the passage number experienced by both our HFLFs and Hayflick’s from the beginning of culture to the dominant senescence was almost the same (~50). However, our culture duration (~0.8 months) from the beginning of culture to the dominant senescence was much shorter than that of Hayflick (~5–6 months). Moreover, based on the theory of telomere shortening with every PD (21), the number of telomere shortening events in the cells of our experiment was much less than that of Hayflick, theoretically. These data indicated that rounding as a source event played a key role in senescence in differentiated normal somatic cells in vitro. This hypothesis is further supported by our experiments showing that the broken chromosomal segment and aneuploid karyotype (Fig. 4D–G) appeared in the HFLFs after only five passages, implying the occurrence of chromosomal instability, a type of genomic instability. Furthermore, our experiments showed that the chromosomal heterogeneity of the HFLFs appeared within the short senescent course (Fig. 4D–G). These data can also explain why telomerase-expressing mouse embryo cells with longer telomeres still exhibit cellular crises (22, 23) similar to that of HFLFs in vitro. These results may be essential for understanding tumorigenesis and senescence.

**Fig. 4.**
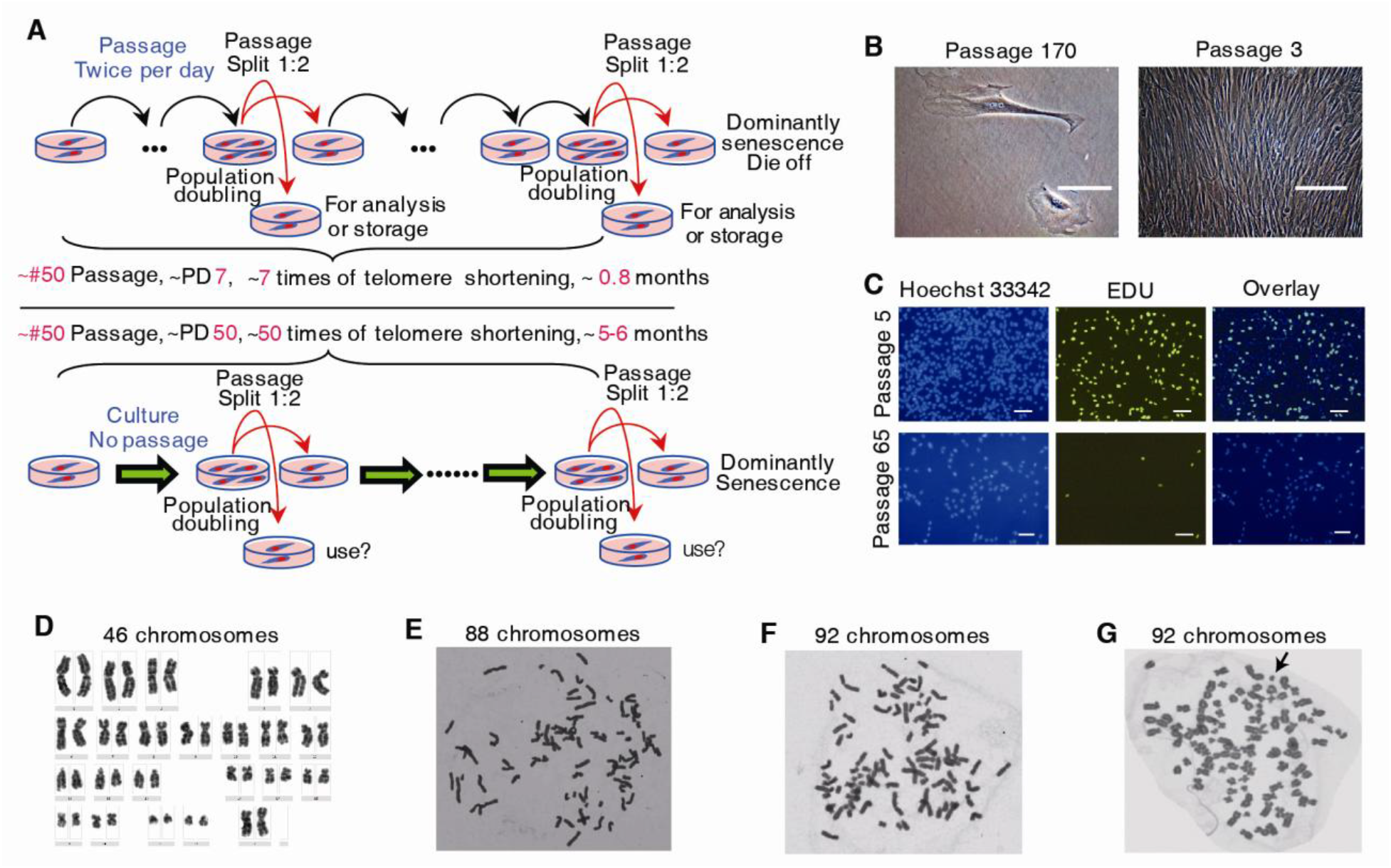
Effects of passage on HFLF senescence and chromosomal instability. (**A**) The procedures of our experiment (upper panel) and the Hayflick-described experiment (lower panel) for testing HFLF senescence. (**B**) Optical images of the morphology of senescent (left) and the young (right) HFLFs. Scale bars, 50 and 100 μm, respectively. (**C**) DNA synthesis experiments with EDU for young HFLFs (five passages; upper row) and senescent cells (65 passages; lower row). Scale bar, 50 μm. (**D**) Normal karyotype of HFLFs after two passages. (**E–G**) Karyotypes of HFLFs after five passages, showing the heterogeneity. The arrow indicates a piece of broken debris in the chromosomes.

Taken together, our results demonstrated the machinery of MIGI, in which cell rounding acts as the initiating event, microtubules function as the driving and switching gear, and SSBs act as the modulating factor. Thus, the components of this machinery suggest that the nonphysiological cell rounding ubiquitously involved in various inflammations (8) in vivo and in cell culture in vitro, including the processes of cell differentiation and expansion for stem cell therapy, may be an important cause of genomic instability and cell senescence for normal somatic cells. Additionally, the experimental data for aneuploidy and DNA shattering may also provide clues for elucidating the mechanisms of heterogeneity, chromothripsis, and kataegis (24-28), which are related to cancer progression, drug resistance, metastasis, recurrence, and patient prognosis. In contrast to normal somatic cells, cancer and embryonic stem cells may avoid catastrophic cell senescence through enlargement of nuclear size, which is a hallmark of these cells (24). These results have important implications for clinical practice. For example, the family of microtubule-targeting chemicals, such as VCR, taxel, and their derivatives, which are routinely used as anticancer drugs in clinical chemotherapy, may become factors for cancer cell’s escaping from deadly MIGI; this feature may further facilitate the optimization of chemotherapy strategies. In addition, for the safety of stem cell therapy (29, 30), common passage methods for cell differentiation and expansion need to be improved so as to reduce the risk of MIGI. Furthermore, understanding the machinery is helpful for revealing the mechanisms underlying some biological activities, such as tumorigenesis and retroviral mutations, which may be involved in the MIGI. In addition, the CISE as a nondestructive detection method may also be applied to investigate the binding states of other DNA-binding partners (see the step by step procedures in the Supplemental Materials and Methods).

## Supplementary Materials

### Materials and Methods

All of the experiments were approved by the ethics committee of our hospital.

#### Isolation of human fetal lung fibroblasts (HFLFs)

The isolation of HFLFs was performed as described (19). The lung tissue was from an aborted female fetus of three month old.

#### Cell culture and maintenance

HFLF, MEF (Zhongke Zesheng Biotechnology Company, Beijing) and HeLa (ATCC) cells were cultured with DMEM (Invitrogen) containing 10% fetal bovine serum (Invitrogen), at the atmosphere of 5% CO2 and 100% humidity. mESCs (E14, ATCC) were cultured under the condition as described (31). All cell passages were performed with 0.25% trypsin.

#### Cell in situ electrophoresis (CISE)

1. 2×10^7^ cells to be analyzed by CISE were acquired by the approach of canonical trypsinization described above.
2. After one time of wash with PBS, the pellet of the cells was diluted with 200μl PBS, and divided into two aliquots putting in two 3.5 cm culture dishes, respectively. Then, the cells in the dishes were distributed evenly on the dish bottom with a pipette tip.
3. The dishes were put on ice and irradiated with 254 nm UV light at the intensity of 1.84W/cm^2^ and total dosage of 165.6J/cm^2^.
4. By flushing the dishes with 20 ml PBS, the irradiated cells were pooled into a 50 ml tube thoroughly.
5. After centrifugation at 260g for 5 minutes and discarding the supernatant liquid, the pellet was gently votexed and prefixed with 200μl of 4% paraformaldehyde for 3 seconds. The prefixation was stopped by quickly adding 46 ml PBS into the tube.
6. After centrifugation at 260g for 5 minutes and discarding the supernatant liquid, the pellet was permeabilized with 200μl of 0.2% Triton X-100 in PBS for 10 minutes. The cells were washed by adding 46 ml of PBS, and centrifuged at 470g for 5 minutes.
7. The cell pellet was diluted with 936μl of CISE buffer (Tris: 25 mM; Glycine: 192 mM; Glucose: 43.2 mM), and gently suspended.
8. 500μl of the cell suspension was pipetted into another 50 ml tube as the control of CISE waiting for the following fixation. The remaining suspension was used for the CISE sample.
9. Assembly the CISE apparatus.
10. Add CISE buffer into the apparatus as the volume settings.
11. Switch on the power, select the mode of constant volts, adjust settings at 500V and run.
12. Puppete 126μl of the CISE sample, and gently drop it into the cell electrophoretic slot at a distance of 1.5 cm from the filter near the cathode.
13. Begin to time the electrophoresis. After running of 3 minutes, stop the electrophoresis.
14. Puppet the cell suspension from the cell electrophoretic slot, and pool it in a 50 ml tube.
15. Thoroughly flush the cell electrophoretic slot with the CISE buffer, and refill the cell electrophoresis slot with the CISE buffer as step 10.
16. Repeat step 11 to 15 until the CISE sample was treated completely.
17. Add CISE buffer to the pooling and control tubes until the liquid volume reaches 50 ml, respectively.
18. Centrifuge the two tubes at 1275g for 5 minutes.
19. Discard the supernatant liquid, and add 300μl of PBS into the two tubes and gently suspend the pellets, respectively.
20. Fix the cells of the two tubes with 15 ml of 70% alcohol (cold), respectively.
21. After fixation of 24 hrs, the cells were tested by flow cytometry.

#### Flow cytometry (FCM) analysis

The fixed cells were washed twice with PBS. Then, the cells with and without CISE were equally divided into four aliquots, respectively. They were used as samples for examination of RPA70, RPA32, POT1, and isotype, respectively. A rabbit polyclonal antibody against RPA70 (A300-241A, Bethyl Labratoryes, Inc.), a rabbit polyclonal antibody against RPA32 (sc-28709, Santa Cruz Biotechnology, Inc.), and a rabbit polyclonal antibody against POT1 (ab21382, Abcam Inc.) were used as primary antibodies for the FCM analyses of all experimental cells. A goat polyclonal FITC-conjugated antibody against rabbit Ig (554020, BD Pharmingen) was used as the secondary antibody for the FCM detection. The incubation concentration of each antibody was adjusted according the recommendation of manufacturer and experimental test. FCM examination was performed with FACSCalibur (Becton, Dickinson Immunocytometry Systems, San Jose, CA). Cell Quest software was used for list mode acquisition, and FlowJo, for analysis. Propidium iodide was used for DNA staining and monitoring cell state. Doublet discrimination mode was used to exclude the aggregation and debris of the cells.

#### Examination of cell population doubling and passage

The treatment of HFLFs, which reached to the doublings, was according to the Hayflick’s methods^8^ except that the cells were detached twice a day with an interval of 6-8 hrs. The procedure of rounding was the same as that of cell passage. When the cells could not reach the confluent growth covering the culture dish bottom, the completed PD and passage numbers were recorded as the cells’ dominant senescence point.

#### Drug treatment of the cells

Vincristine (Zhejiang Hisun pharmaceutical Co., Ltd) were used to modulate the assembly and disassembly of microtubules in the HFLFs, MEFs, HeLa cells and mESCs, respectively. The final concentration of Vincristine in the medium is 100ng/ml. The treatment time with Vincristine was 1.5hrs.

#### Cell proliferation examination

The proliferation of HFLFs at passage 5 and passage 65 were examined, respectively, with Click-iT^®^ EDU Imaging Kits (C10337, Invitrogen). The procedure was according to the instruction manual of the Kits. The EDU labeling time was 1 hr.

#### Nuclear staining and microscopic imaging

For staining spreading cells, the experimentally treated cells growing in a 3-cm dish, at exponential phase, were washed with PBS for two times. Then, the cells were covered with 1 ml of Hoechst 33342 solution (Invitrogen, 5μg/ml in PBS) for 30 min, at room temperature, in protection from light. After two washes with PBS, the cells were examined under a Nikon TE300 microscope. For staining rounded cells, the experimentally treated cells growing in a 3-cm dish, at exponential phase, were collected in a 1-ml tube, by the canonical trypsinization. Then, the cells were stained with 1 ml of Hoechst 33342 solution for 30 min. After two washes with PBS, the cells were suspended with 200μl of PBS and microscopically examined by dropping the cells on slides.

#### Karyotyping analysis

Karyotypes of HFLFs with passage 2 and 5 were analyzed as described previously (32). Nocodazole was used for mitotic arrest. The final concentration is 0.1 μg/ml. And the incubation time is 1.5 hrs.

## References

1. D. J. Richard, E. Bolderson, K. K. Khanna, Multiple human single-stranded DNA binding proteins function in genome maintenance: structural, biochemical and functional analysis. Crit. Rev. Biochem. Mol. Biol. 44, 98-116 (2009).

2. T. Veldman, K. T. Etheridge, C. M. Counter, Loss of hPot1 function leads to telomere instability and a cut-like phenotype. Curr. Biol. 14, 2264-2270 (2004).

3. P. Baumann, T. R. Cech, Pot1, the putative telomere end-binding protein in fission yeast and humans. Science 292, 1171-1175 (2001).

4. L. Zou, S. J. Elledge, Sensing DNA damage through ATRIP recognition of RPA-ssDNA complexes. Science 300, 1542-1548 (2003).

5. H. Chen, M. Lisby, L. S. Symington, RPA coordinates DNA end resection and prevents formation of DNA hairpins. Mol. Cell 50, 589-600 (2013).

6. M. Dobbelstein, C. S. Sørensen, Exploiting replicative stress to treat cancer. Nat. Rev. Drug. Discov. 14, 405-423 (2015).

7. J. T. Yeeles, T. D. Deegan, A. Janska, A. Early, J. F. Diffley, Regulated eukaryotic DNA replication origin firing with purified proteins. Nature 519, 431-435 (2015).

8. J. Berkes, V. K. Viswanathan, S. D. Savkovic, G. Hecht, Intestinal epithelial responses to enteric pathogens: effects on the tight junction barrier, ion transport, and inflammation. Gut 52, 439-451 (2003).

9. J. R. Sims, S. Karp, D. E. Ingber, Altering the cellular mechanical force balance results in integrated changes in cell, cytoskeletal and nuclear shape. J. Cell Sci. 103, 1215-1222 (1992).

10. R. L. Creager, Y. Li, D. M. MacAlpine, SnapShot: Origins of DNA replication. Cell 161, 418-418 (2015).

11. K. H. Chow, R. E. Factor, K. S. Ullman, The nuclear envelope environment and its cancer connections. Nat. Rev. Cancer 12, 196-209 (2012).

12. K. N. Dahl, A. J. Ribeiro, J. Lammerding, Nuclear shape, mechanics, and mechanotransduction. Circ. Res. 102, 1307-1318 (2008).

13. L. J. Edens, K. H. White, P. Jevtic, X. Li, D. L. Levy, Nuclear size regulation: from single cells to development and disease. Trends Cell Biol. 23, 151-159 (2013).

14. A. Downes, R. Mouras, A. Elfick, Optical spectroscopy for noninvasive monitoring of stem cell differentiation. J. Biomed. Biotechnol. 2010, 1-10 (February 2010).

15. X. Wang et al., MAD2-induced sensitization to vincristine is associated with mitotic arrest and Raf/Bcl-2 phosphorylation in nasopharyngeal carcinoma cells. Oncogene 22, 109-116 (2003).

16. A. Mazumder et al., Prestressed nuclear organization in living cells. Methods Cell Biol. 98, 221-239 (2010).

17. A. Mazumder, G. V. Shivashankar, Emergence of a prestressed eukaryotic nucleus during cellular differentiation and development. J. R. Soc. Interface 7 Suppl 3, S321-330 (2010).

18. A. Adey et al., The haplotype-resolved genome and epigenome of the aneuploid HeLa cancer cell line. Nature 500, 207-211 (2013).

19. L. Hayflick, The limited in vitro lifetime of human diploid cell strains. Exp. Cell Res. 37, 614-636 (1965).

20. M. Wadman, Medical research: cell division. Nature 498, 422-426 (2013).

21. C. B. Harley, A. B. Futcher, C. W. Greider, Telomeres shorten during ageing of human fibroblasts. Nature 345, 458-460 (1990).

22. T. A. Carman, C. A. Afshari, Barrett, J. C. Cellular senescence in telomerase-expressing Syrian hamster embryo cells. Exp. Cell Res. 244, 33-42 (1998).

23. N. E. Sharpless, C. J. Sherr, Forging a signature of in vivo senescence. Nat Rev Cancer 15, 397-408 (2015).

24. N. McGranahan, C. Swanton, Biological and therapeutic impact of intratumor heterogeneity in cancer evolution. Cancer Cell 27, 15-26 (2015).

25. I. Martincorena, P. J. Campbell, Somatic mutation in cancer and normal cells. Science 349, 1483-1489 (2015).

26. H. Gaillard, T. García-Muse, A. Aguilera, Replication stress and cancer. Nat. Rev. Cancer 15, 276-289 (2015).

27. M. Kansara, M. W. Teng, M. J. Smyth, D. M. Thomas, Translational biology of osteosarcoma. Nat. Rev. Cancer 14, 722-735 (2014).

28. A. Aguilera, T. García-Muse, Causes of genome instability. Annu. Rev. Genet. 47, 1-32 (2013).

29. I. J. Fox et al., Use of differentiated pluripotent stem cells as replacement therapy for treating disease. Science 345, 1247391 (2014).

30. A. S. Lee, C. Tang, M. S. Rao, I. L. Weissman, J. C. Wu, Tumorigenicity as a clinical hurdle for pluripotent stem cell therapies. Nat. Med. 19, 998-1004 (2013).

## References (31-32)

31. M. Takenaga, M. Fukumoto, Y. Hori, Regulated Nodal signaling promotes differentiation of the definitive endoderm and mesoderm from ES cells. J. Cell Sci. 120, 2078-2090 (2007).

32. D. Moralli et al., An improved technique for chromosomal analysis of human ES and iPS cells. Stem Cell Rev. 7, 471-477 (2011).

